# Genetically modified primary airway-epithelial cell culture models to study virus-host interactions of emerging respiratory viruses

**DOI:** 10.64898/2026.10.16.758770

**Authors:** Karen Stevers, Sarah Jordi, Blandina I. Oliveira Esteves, Ronald Dijkman

## Abstract

Airway epithelial cell (AEC) cultures such as air-liquid interface cultures or airway organoids are widely used to study emerging respiratory virus infection in a physiologically representative model system of the human respiratory epithelium. Loss-of-function studies in AEC culture systems offer a unique opportunity to better understand viral disease mechanisms of emerging viruses in the human host, may reveal biologically relevant host-factor dependencies that can be therapeutically targeted, or identify human host barriers of cross-species transmission of emerging viruses that can inform risk assessment. In this chapter, we describe the generation of knockout (KO) AECs using engineered virus-like particles (eVLPs). These particles transiently deliver the Cas9–gRNA ribonucleoprotein (RNP) complex into target cells, enabling efficient genome editing of any gene of interest. We detail the production, quantification, and application of eVLPs for gene editing in AECs, alongside an optimized protocol for AEC expansion and differentiation at the air-liquid interface that is compatible with genetically modified cells. Finally, we describe procedures for infecting KO AECs with emerging respiratory viruses. Together, these protocols provide a robust workflow for generating renewable KO AEC cultures in less than two months, facilitating the functional characterization of host factors that influence viral infection of the airway epithelium.

## 1. Introduction

Emergence of novel viral pathogens in the human population represents a major global health threat, and can lead to global pandemics due to a lack of pre-existing immunity [1]. Most of these pathogens are of animal origin, and emerging viruses must adapt to effectively exploit a new host for replication and spread [2]. Adaptation occurs at many levels of the viral replication cycle, including gaining the ability to bind and use host cell receptors for entry, replicating its genome using host cell machinery, and overcoming host restriction factors such as the antiviral innate immune response [3].

Studying the role of human host factors during infection with emerging viruses in human cells is crucial for understanding mechanisms of disease pathogenesis and might reveal host-factor dependencies that can be therapeutically targeted or identify human host barriers to cross-species transmission that can inform risk assessment. Functional analysis of host factors is commonly carried out by loss- or gain-of-function studies in transgenic small animal models or in immortalized cell lines. However, suboptimal representation of these model systems to infection in human tissues may lead to discoveries that are challenging to translate to meaningful clinical interventions.

The airway epithelium is the primary target of numerous respiratory viruses, including (emerging) coronaviruses and influenza A viruses. Human airway epithelial cell (AEC) culture models, such as airway organoids (AOs) and air-liquid interface (ALI) cultures, closely recapitulate the cellular diversity, architecture, and physiological functions of the human airway epithelium under controlled *in vitro* conditions [4–6]. Loss-of-function studies in AEC culture systems therefore offer a unique opportunity to identify biologically relevant host factors that regulate infection by emerging respiratory viruses in the airway epithelium.

Cells in which one specific gene is inactivated, known as knockout (KO) cells, are usually generated by programmable nucleases such as Cas9 that are targeted to specific sites in the genome by a guide RNA (gRNA) [7]. Cas9 nucleases and corresponding gRNAs can be delivered to target cells in various formats via diverse methods, including viral vectors, electroporation, and lipofection. Our group and others have previously demonstrated that viral delivery is particularly efficient for targeting AECs [8, 9]. However, conventional lentiviral vectors permanently integrate Cas9 and the gRNA into the host genome, raising concerns about genome instability due to random integration and prolonged Cas9 expression [10]. To address these limitations, we recently evaluated a transient viral delivery system for gene editing in primary airway epithelial cells (Stevers et al, 2026, manuscript in preparation), called engineered virus-like particles (eVLPs). eVLPs are assemblies of viral proteins that lack a viral genome, but can exploit viral assembly mechanisms to package Cas9-gRNA ribonucleoproteins (RNPs) into virions [10]. They also retain the cell entry properties of enveloped viruses, enabling efficient delivery of RNP complexes to target cells, where the RNP complex mediates genome editing with minimal off-target effects owing to its short lifetime [11, 12]. Using this technology, we successfully disrupted the *MX1* gene in AECs and investigated its role in the interferon-mediated antiviral response to influenza A virus infection (Stevers et al, 2026, manuscript in preparation).

In this chapter, we describe procedures for producing and quantifying engineered virus-like particles (eVLPs), as well as their application for gene editing in AECs. We also present an updated protocol for expanding AECs and differentiating them at the air-liquid interface (ALI), compatible with genetically modified cells. This protocol builds on our previously published methods for the establishment and infection of primary airway epithelial cell models [13, 14]. A key improvement is the incorporation of airway organoid expansion protocols, which enable efficient amplification of primary airway epithelial cells while preserving their stem cell characteristics and extending their lifespan compared with conventional two-dimensional culture methods [4]. Together, these protocols provide a robust workflow for generating renewable KO AEC cultures in under two months (Figure 1) and facilitate the functional characterization of host factors involved in emerging viral infections of the airway epithelium.

**Figure 1:**
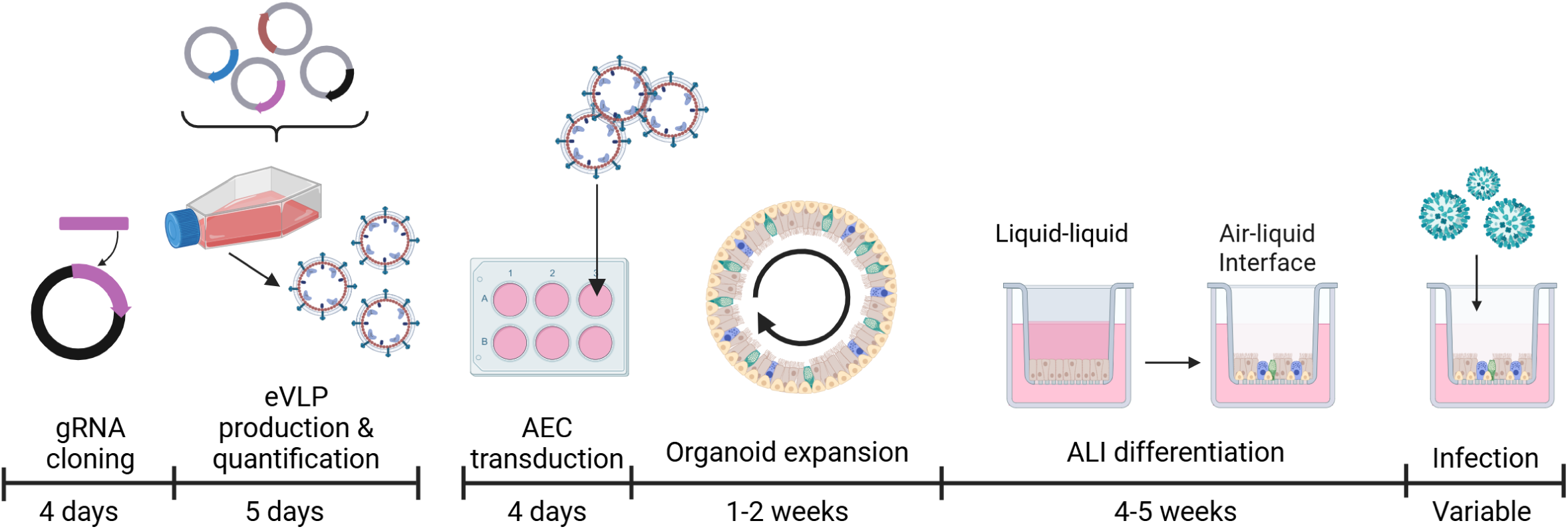
Chronological representation of the generation of KO AECs. A gRNA targeting a gene of interest is cloned into the SUPERBLADE5 vector. Corresponding eVLPs are produced by transducing 293 LTV cells with plasmids encoding the gRNA, a VSV glycoprotein, Cas9, and structural proteins to form the virion, and are quantified. Subsequently, AECs are transduced with eVLPs, expanded into organoids, and and subcultured onto porous support chambers. After an initial expansion in a liquid-liquid phase, AECs are differentiated at the air-liquid interface for at least 4 weeks. Afterwards, KO AECs are ready to be infected with emerging respiratory viruses. Created in BioRender. Stevers, K. M. (2026) https://BioRender.com/jg8hqg3.

## 2. Materials

### 2.1 eVLP production

#### 2.1.1 gRNA design and cloning

1. 10x T4 ligase buffer (Promega)
2. SUPERBLADE5 plasmid (Addgene #134913)
3. BsmBI-V2 restriction enzyme (New England Biolabs, 10,000 units/mL)
4. 0.5% agarose gel in Tris-Acetate EDTA buffer with 1x RedSafe (Intron Biotechnology) + agarose gel electrophoresis equipment
5. E.Z.N.A gel extraction kit (Omega Bio-tek), or equivalent
6. T4 Ligase (Promega)
7. Stellar chemically competent bacteria (or equivalent) and SOC medium (Takara Bio)
8. Luria broth (LB) agar plates with 50 µg/mL Carbenicillin
9. LB medium with 50 µg/mL Carbenicillin
10. Nucleobond Xtra Midiprep kit (Macherey-Nagel) or equivalent plasmid DNA extraction kit
11. M13 primer: 5’-TGTAAAACGACGGCCAG-3’

#### 2.1.2 eVLP production in 293-LTV cells

1. Poly-D-Lysine (Gibco), 1:1 diluted in Sterile Phosphate-Buffered Saline (PBS) to a final concentration of 50 µg/mL.
2. 6-well cell culture plate or T75 or T175 cm2 flasks (tissue-culture treated)
3. Sterile ultrapure water
4. 293-LTV cells (Cell Biolabs)
5. 293-LTV medium: Dulbecco’s Modified Eagle Medium, high glucose (DMEM), supplemented with 2 mM GlutaMax (Gibco), 10% fetal bovine serum (Pan Biotech), 100 U/mL penicillin, 100 μg/mL streptomycin (Gibco), 1x MEM Non-Essential Amino Acids (Gibco) and 15 mM HEPES (Gibco)
6. JetOptimus transfection reagent and JetOptimus buffer (Polyplus)
7. SUPERBLADE5_gRNA plasmid, from section 2.1.1
8. pCMV-MMLVgag-3xNES-Cas9 plasmid (Addgene #181752)
9. pBS-CMV-gagpol plasmid (Addgene #35614)
10. pMD2.G plasmid (Addgene #12259)
11. Sterile PBS
12. Complete Pneumacult Ex Plus medium (Stemcell): PneumaCult-Ex Plus Basal Medium supplemented with 1x PneumaCult Ex Plus 50x Supplement, 0.48 µg/mL Hydrocortisone (Stemcell) and 100 μg/mL Primocin (Invivogen). Complete Ex Plus medium can be stored at 4°C for 4 weeks.
13. Epithelial expansion medium (EpiX): Supplement complete Pneumacult Ex Plus medium directly before use with EpiX additives, 1 µM A-83-01, 5 µM Y-27632, 3 µM Isoprotenerol hydrochloride. Prepare a stock of A-83-01 (10 mM, 1:10,000) by dissolving 10 mg in 2.37 mL DMSO. Prepare a stock of isoprotenerol hydrochloride (100 mM) by dissolving 100 mg isoprotenerol in 4 mL sterile ultrapure water. Further dilute in ultrapure water to 30 mM (1:10,000 stock). Prepare a Y-27632 stock (20 mM, 1:4000) by dissolving 10 mg Y-27632 in 1.56 mL sterile ultrapure water, protect from light. Additives should be stored at -20°C for up to 1 year.

#### 2.1.3 eVLP concentration

1. 15 mL Amicon® ultra centrifugal filter (100 kDa molecular weight cut-off)
2. Bovine serum albumin (BSA) blocking solution: Dissolve 1 g cell culture grade BSA (Jackson Immunoresearch) in 100 mL PBS to get a 1% BSA solution. Filter-sterilize the solution with a 0.22 µm filter.
3. Sterile ultrapure water
4. EpiX medium
5. Optional: Pluronic F-68 (Gibco)

#### 2.1.4 eVLP quantification

1. 5% Triton X-100 in PBS: Dilute 5 mL Triton® X-100 in 95 mL PBS
2. Cas9 ELISA kit (Cell Biolabs)
3. Spectrophotometer (450 nm)

### 2.2 AEC transduction

#### 2.2.1 collagen coating of cell culture ware

1. Collagen solution from human fibroblasts (Sigma-Aldrich, Type I, 3 mg/mL)
2. Sterile ultrapure water
3. Sterile PBS
4. 24-well cell culture plate

#### 2.2.2 Thawing and seeding of primary airway epithelial cells with CEPT

1. Primary airway epithelial cells, see note 1.
2. Thawing medium: Complete Pneumacult Ex Plus medium freshly supplemented with 3 µM Isoprotenerol hydrochloride, 1 µM A83-01 and 1x CEPT (50 nM Chroman 1, 5 µM Emricasan, 700 nM trans-ISRIB and 1x Polyamine, see note 2).
3. Collagen-coated cell culture plate

#### 2.2.3 Transduction of AECs with eVLPs

1. Concentrated eVLPs
2. EpiX medium
3. Polybrene (Sigma-Aldrich, 10 mg/mL)
4. Hanks Balanced Salt Solution (Gibco, HBSS)

#### 2.2.4 Evaluating gene editing efficiency with ICE

1. eVLP-treated and control AECs
2. DirectPCR lysis reagent (Viagen)
3. Proteinase K (Promega, 20 mg/mL)
4. GoTaq G2 Green master mix (Promega), or equivalent polymerase master mix
5. 1.5% agarose gel in Tris-Borate EDTA buffer with 1x RedSafe + agarose gel electrophoresis equipment
6. E.Z.N.A Cycle Pure Kit (Omega Bio-tek), or equivalent PCR purification kit.

### 2.3 KO AEC expansion and differentiation at ALI

#### 2.3.1 Organoid expansion of KO AECs

1. Cultrex™ Reduced Growth Factor Basement Membrane Extract, Type 2 (BME) (R&D systems)
2. 48-well suspension culture plates
3. HBSS
4. TrypLE™ Express Enzyme Reagent (Gibco)
5. Pre-AOM: Advanced DMEM/F12 supplemented with 2 mM Glutamax, 100 U/mL penicillin, 100 μg/mL streptomycin, 10 mM HEPES. Store at 4°C for up to 6 months.
6. (Optional): Trypan blue and counting chambers
7. Airway organoid medium (AOM). Prepare reagents according to ***Table 1***, prepare medium according to ***Table 2***, see note 3.

#### 2.3.2 Collagen Type IV Coating of porous support chambers

1. Porous support chambers (diameter 6.5 mm, pore size 0.4 μm) in 24-well plates.
2. 0.5 mg/mL Collagen type IV from human placenta. Dissolve 10 mg in 5 mL 0.25% acetic acid in ultrapure water at 37°C for several hours. Swirl occasionally until dissolved. Add 0.25% acetic acid in ultrapure water to a total volume of 20 mL. Filter-sterilize the Collagen IV solution through a 0.22 μm filter and store at −20°C for at least 1 year.
3. Sterile ultrapure water
4. Sterile PBS

#### 2.3.3 Seeding, differentiation and maintenance of KO AEC

1. EpiX medium
2. HBSS (Gibco)
3. Dispase II solution (Sigma-Aldrich). Reconstitute 100 mg Dispase II in 10 mL 50 mM Hepes/KOH pH 7.4, 150 mM NaCl solution to 10 mg/mL by pipetting up and down. Aliquot in 1 mL and store at -20°C for at least 1 year. Directly before use, dilute to 1 mg/mL in DMEM supplemented with 100 U/mL penicillin, 100 μg/mL streptomycin, 15 mM HEPES.
4. TrypLE™ Express Enzyme Reagent
5. Pre-AOM supplemented with 5 µM Y-27632
6. TEER solution: dissolve 4.5 g NaCl, 91.89 mg CaCl2, and 1.194 g of HEPES in 400 mL of ultrapure water and stir for 10 min at room temperature. Add water up to 500 mL and sterilize the solution via autoclaving, 20 min at 121°C.
7. ALI medium. First prepare Pneumacult ALI complete base medium (Stemcell) by mixing 450 mL PneumaCult-ALI Basal Medium with 50 mL PneumaCult-ALI 10x Supplement. Aliquot in 40 mL and freeze at -20°C for at least 1 year. Prepare Complete Pneumacult ALI maintenance medium according to **Table 3**. Store at 4°C for up to 2 weeks.

#### 2.3.4 Cryopreservation of KO AEC

1. Cryostor CS10 cryopreservation medium (Stemcell)
2. Cryovials
3. Pre-AOM, supplemented with 1x CEPT
4. BME
5. 48-well suspension culture plates
6. AOM, replace 5 µM Y-27632 with 1x CEPT.

### 2.4 KO AEC

#### 2.4.1 Virus infection of KO AECs

1. TEER solution
2. HBSS
3. ALI medium
4. Virus transport medium (VTM): Modified Eagle Medium (MEM), 25 mM HEPES-buffered, 0.5% gelatin, 100 U/mL penicillin and 100 μg/mL streptomycin

## 3. Method

### 3.1 eVLP production

#### 3.1.1 gRNA design and cloning

To direct Cas9 to a gene of interest, a gRNA targeting this DNA sequence must be designed and cloned in the SUPERBLADE5 expression vector. See also note 4.

1. gRNAs can be designed using webtools such as CHOPCHOP (https://chopchop.cbu.uib.no/) [15] or can be found within CRISPR libraries, such as the Moffat library (https://crispr.ccbr.utoronto.ca/public/library/TKOv3/) [16]. Select for SpCas9.
2. Select gRNAs binding in an exon present in all variants/isoforms of the targeted gene. If possible, select an early exon. See also note 5.
3. Delete the PAM site from your selected sequences and generate the reverse complement
4. Add the overhangs to the sequences for cloning into the SUPERBLADE5 plasmid and order forward and reverse oligonucleotides. For the BsmBI restriction enzyme, this means: add a “G” to the 5’ end of your forward guide sequence if it does not start with a “G”, add “CACC” to the 5’ end of your forward guide sequence, add “AAAC” at the 5’ end of your reverse guide sequence and add “C” at the 3’ end of your reverse guide sequence when the guide sequence did not start with an “G”:

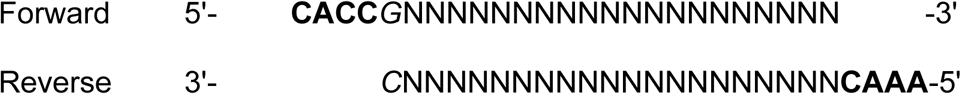
5. Anneal the forward and reverse guide oligos by mixing both oligos with T4 ligase buffer (final concentration 1x) to a final concentration of 10 µM. Incubate the mixture in a thermocycler at 95°C for 5 min, then cool to 25°C at a ramp rate of 0.1°C/s.
6. Digest the SUPERBLADE5 plasmid with the BsmBI-V2 enzyme by mixing 3 µg plasmid with 2 µL of enzyme in the corresponding digestion buffer and incubate at 55°C for 2h.
7. Load the digested plasmid on a 0.5% agarose gel and separate the fragments by size using agarose gel electrophoresis. Cut out the 3.3 kb fragment and perform gel cleanup according to the manufacturer’s instructions.
8. Ligate the annealed gRNA and the digested backbone plasmid by adding 50 ng plasmid, 1 µL double-stranded gRNAs (1:200 diluted), and 1 µL T4 ligase to 1 µL T4 ligase buffer in sterile water (total volume 10 µL), and incubate at RT for 30 min.
9. Transform the plasmid into Stellar competent cells by adding 2 µL of the ligation product to 20-40 µL of bacteria. Incubate on ice for 30 min, then heat-shock the bacteria at 42°C for 40-45 sec. Cool the transformation solution on ice for 2 min, then resuspend in 500 µL of SOC medium. Plate 20 µL of the solution onto LB/carbenicillin agar plates and incubate at 37°C overnight.
10. Pick 2-3 colonies and expand the bacteria in LB/carbenicillin medium.
11. Isolate plasmid DNA using the NucleoBond Xtra Midiprep kit and confirm correct incorporation of the gRNA into the plasmid by Sanger Sequencing. For sequencing, use the M13 forward primer.

**Table 1:** Airway organoid medium additives.

| **Media component** | **Supplier** | **Stock concentration** | **Solvent** | **Remarks** |
| --- | --- | --- | --- | --- |
| R-Spondin 1 | EPFL Protein Expression<br>Core Facility/Peprotech | 100 µg/mL | PBS, 0.1% BSA |  |
| Noggin | EPFL Protein Expression<br>Core Facility/Peprotech | 100 µg/mL | PBS, 0.1% BSA |  |
| Fibroblast growth factor 7 | Peprotech | 100 µg/mL | PBS, 0.1% BSA |  |
| Fibroblast growth factor 10 | Peprotech | 100 µg/mL | 5 mM sodium phosphate, pH 7.4,<br>0.1% BSA |  |
| B27 supplement | Gibco | 50x | NA |  |
| N-Acetylcysteine | Sigma-Aldrich | 500 mM | H2O | Filter-sterilize through a 0.22 µm filter |
| Nicotinamide | Sigma-Aldrich | 1M | PBS | Filter-sterilize through a 0.22 µm filter |
| SB202190 | Sigma-Aldrich | 5 mM | DMSO | Protect from light |
| Y-27632 | Tocris | 20 mM | H2O | Protect from light |
| A83-01 | Tocris | 10 mM | DMSO |  |

**Table 2:** Airway organoid medium composition.

| Media component | Volume for 10 mL AOM | Final concentration |
| --- | --- | --- |
| Pre-AOM | 9.66 mL | NA |
| R-Spondin 1 | 50 µL | 500 ng/mL |
| Noggin | 10 µL | 100 ng/mL |
| Fibroblast growth factor 7 | 2.5 µL | 25 ng/mL |
| Fibroblast growth factor 10 | 10 µL | 100 ng/mL |
| B27 supplement | 200 µL | 1x |
| N-Acetylcysteine | 25 µL | 1.25 mM |
| Nicotinamide | 50 µL | 5 mM |
| SB202190 | 1 µL | 500 nM |
| Y-27632 | 2.5 µL | 5 mM |
| A83-01 | 0.5 µL | 500 nM |

**Table 3:** ALI medium composition.

| Media component | Volume | Stock concentration | Final concentration | Supplier |
| --- | --- | --- | --- | --- |
| Pneumacult ALI complete base medium | 40 mL |  |  | StemCell |
| PneumaCult-ALI Maintenance Supplement (100×) | 400 µL | 100x | 1x | StemCell |
| Primocin | 80 µL | 50 mg/mL | 100 µg/mL | Invivogen |
| Hydrocortisone | 200 µL | 96 µg/mL | 0.48 µg/mL | StemCell |
| Heparin | 80 µL | 2 mg/mL | 4 µg/mL | StemCell |

#### 3.1.2 eVLP production in 293 LTV cells

Produce eVLPs by transfecting 293 LTV cells. See note 6.

1. Coat a cell culture flask with Poly-D-Lysine (50 µg/mL, 10 mL for T175, 5 mL for T75 flask, 1 mL per 6-well) and incubate it at 37°C in a humidified incubator with 5% CO_2_ for 2 h. Aspirate and wash the vessel three times with sterile ultrapure water.
2. Seed 293-LTV cells in coated cell culture vessel in LTV medium one day prior to transfection (for cell density, see
3. ***Table 4***). Incubate in a humidified incubator at 37°C with 5% CO₂.
4. Prepare the transfection mix by adding the plasmid DNA to the JetOptimus buffer (for plasmid/transfection reagent ratios, see
5. ***Table 4***). Vortex the mixture and add JetOptimus transfection reagent, vortex and incubate at RT for 10 min.
6. Add the transfection mix dropwise to the cells and incubate at 37°C in a humidified 5% CO_2_ incubator for 24h.
7. Aspirate the medium, wash twice with PBS and replace with EpiX medium.
8. Incubate the cells in a humidified incubator at 37°C with 5% CO₂ until 72 h post-transfection.
9. Collect the eVLP-containing supernatant from the 293-LTV cells and collect it in a 50 mL conical tube. Clear the supernatant by centrifugation at 1000 x g for 5 min at RT. Carefully transfer the supernatant to a fresh 50 mL conical tube. For unconcentrated use, aliquot and store at -80°C (see note 6). For concentrated use, continue with step 3.1.3.

**Table 4:** eVLP production in 293-LTV cells in different formats.

| Component | 6-well | T75 | T175 |
| --- | --- | --- | --- |
| Seeding density 293-LTV cells | 8.8 E5 | 6.6 E6 | 15.4 E6 |
| **Plasmids** |  |  |  |
| pBS_CMV-gagpol | 753 ng | 6,033 ng | 13,849 ng |
| pCMV-MMLVgag-3xNES-Cas9 | 457 ng | 3,656 ng | 8,409 ng |
| pMD2.G | 108 ng | 860 ng | 1,978 ng |
| SUPERBLADE5_gRNA | 1,183 ng | 9,462 ng | 21,763 ng |
| JetOPTIMUS buffer | to 200 µL | to 1500 µL | to 3500 µL |
| JetOPTIMUS reagent | 2.5 µL | 20 µL | 46 µL |

#### 3.1.3 eVLP concentration

eVLPs are concentrated by ultrafiltration to increase dose and remove contaminants derived from producer cells.

1. Wash Amicon Ultra centrifugal filter tubes with 15 mL of sterile ultrapure water, then centrifuge at 1500 × g for 5 min at RT. Remove the remaining water by pipetting, add 15 mL of BSA blocking solution, and incubate for at least 2 h at RT. Discard blocking solution and rinse the filter 3x with ultrapure water. Centrifuge at 1500 × g for 5 min at RT, then aspirate the remaining water.
2. Add up to 15 mL of eVLP supernatant to the blocked Amicon filter tube, then centrifuge at 1500 x g at 4°C until the eVLPs are 25-50x concentrated. 15 mL solutions take approximately 45-60 min of centrifugation.
3. Collect the concentrate and store it in a tube at 4°C.
4. Add the remaining eVLP supernatant and repeat the concentration until the complete supernatant is concentrated, see note 7.
5. Wash the filter with 200-300 µL of EpiX medium to recover the eVLPs by pipetting the medium onto the filter and collecting it at the bottom. Pipet tips can lightly touch the filter, but the filter should not be scratched. Repeat washing 5x per side.
6. Add the recovered eVLPs to the already collected concentrate and mix well. Optional: Add Pluronic F-68 (1:100 diluted) to prevent eVLPs from sticking to the plastic.
7. Aliquot the eVLPs and store at -80°C. Use within 3 months.

#### 3.1.4 eVLP quantification

The eVLPs are quantified by measuring the amount of Cas9 protein in the eVLP prep using a commercially available Cas9 ELISA kit. The eVLPs are inactivated with Triton X-100 before performing the ELISA.

1. Dilute the eVLPs in assay diluent. For 50x concentrated eVLPs, use a 1:250 and 1:1000 dilution, or adjust accordingly.
2. Transfer 225 µL of each sample to a microcentrifuge tube containing 25 µL of 5% Triton X-100 solution.
3. Dilute the standard (5 µg/mL) to 100 ng/mL by diluting 10 µL in 440 µL assay diluent and 50 µL 5% triton X-100. Prepare a 7-step two-fold dilution series from 100-1.56 ng/mL Cas9 by diluting 250 µL of the previous concentration with 225 µL assay diluent and 25 µL Triton X-100. Prepare a blank by mixing 225 µL assay diluent with 25 µL Triton X-100.
4. Vortex the samples, standards, and blank, and incubate at 37°C for 30 min.
5. Load samples and standards in duplicates (2 × 100 µL) to the ELISA plate. Follow the manufacturer’s instructions for the remainder of the ELISA protocol.
6. Read absorbance of each sample and standard using a spectrophotometer at 450 nm. Define a function for the standard curve between absorbance and Cas9 concentration and extrapolate the concentration for each eVLP prep.

### 3.2 AEC transduction

#### 3.2.1 Collagen coating of cell culture ware

Cell culture ware should be coated with collagen type I to ensure proper attachment of AECs to the plastic.

1. Dilute the collagen solution to 0.04 mg/mL in sterile ultrapure water.
2. Add 250 µL per 24-well, or adjust volume for area accordingly.
3. Incubate for 2 - 4 h in a humidified incubator at 37°C with 5% CO₂.
4. Aspirate the remaining coating solution, leave 5 min to airdry, and wash twice with PBS.
5. Use immediately or store at 4°C for up to 6 weeks.

#### 3.2.2 Thawing and seeding of primary airway epithelial cells with CEPT

Primary airway epithelial cells are thawed in EpiX medium supplemented with CEPT cocktail to improve viability. The CEPT cocktail consists of Chroman 1, Emricasan, Polyamine and trans-ISRIB and reduces cell stress during cryopreservation, thawing and passaging [17].

1. To thaw cryopreserved cells, dip a cryovial in a water bath (20-37°C) for 1-2 min and transfer the contents to a 15 mL conical tube. Add 1 mL of prewarmed (37°C) EpiX medium supplemented with CEPT dropwise at a rate of one drop every 2 sec. Then slowly and dropwise add an additional 3 mL of prewarmed EpiX supplemented with CEPT at a rate of two drops every second (up to 5mL). Mix slowly by pipetting up and down.
2. Centrifuge the cells at 200 x g for 3 min at RT and carefully remove the supernatant
3. Resuspend the cells in 1 mL EpiX supplemented with CEPT and count the cells with Trypan blue.
4. Resuspend the cells to a density of 100,000 cells/mL in EpiX medium supplemented with CEPT, and seed 500 µL at 50,000 cells per well in each well of a collagen-coated 24-well plate (or adjust volume/size accordingly).
5. Culture cells overnight in a humidified incubator at 37°C with 5% CO₂.

#### 3.2.3 Transduction of AECs with eVLPs

During transduction, eVLPs complexed with non-targeting gRNAs can be used as controls; see note 5. An example of cell morphology and density of AECs before and after eVLP transduction can be found in Figure 2.

1. Thaw eVLPs on ice.
2. Dilute eVLPs in EpiX medium to a final concentration of 0.2-0.4 µg/mL (based on Cas9, quantified by ELISA). See also note 8.
3. Add Polybrene (final concentration 5 µg/mL) to the eVLP dilution.
4. Aspirate medium from the cells and add the eVLP/polybrene/medium mix (500 µL in a 24-well, 2.5 mL in a 6-well)
5. Incubate cells in a humidified incubator at 37°C with 5% CO₂ for 6h-8h. Then aspirate the inoculum, wash the cells with HBSS, and add fresh EpiX medium.
6. Incubate the cells in a humidified incubator at 37°C with 5% CO₂ until 72 h post-transduction.
7. Continue with cell harvest for genome editing efficiency analysis or cell expansion.

**Figure 2:**
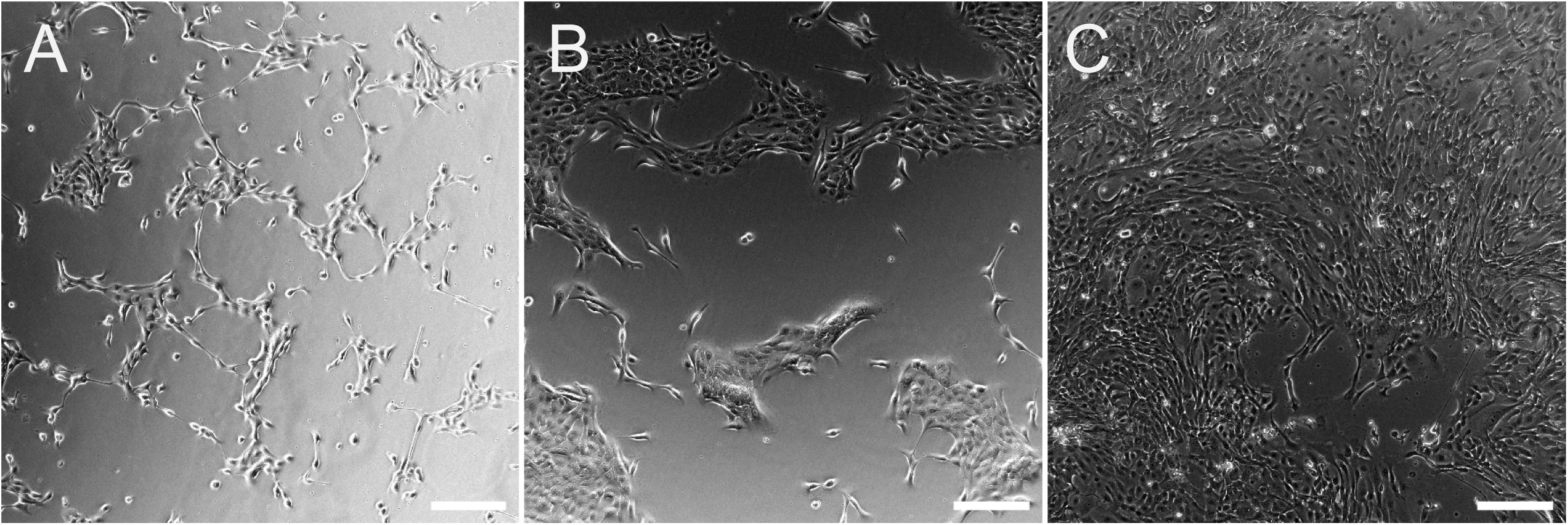
Morphology of AECs pre- and post-transduction with eVLPs. Representative brightfield images of AECs 24 h post-seeding, directly before transduction **(A),** 24 h post-transduction **(B)** and 72 h post-transduction, before harvest **(C).** Scale bar = 200 µm.

#### 3.2.4 Evaluating gene editing efficiency with ICE

To evaluate gene-editing efficiency at the genome level, isolate DNA, amplify and sequence the targeted locus, and analyze efficiency using the online tool ICE (Inference of CRISPR Edits, [18]). To confirm the knockout at the protein or functional level, perform further analyses such as flow cytometry, western blot, or ELISA, following established protocols not described in this chapter.

An example of results that can be expected by following this method, using a gRNA targeting the *STAT1* gene, can be found in Figure 3.

1. Design primers to amplify the region 200 bp upstream and 500 bp downstream of the gRNA target site in your gene of interest.
2. Prepare cell lysis buffer by diluting proteinase K to 200 µg/mL in DirectPCR lysis reagent.
3. Lyse cells in lysis buffer (1000 cells/µL), by either adding lysis buffer directly on cell monolayer or by resuspending a cell pellet collected at any point 72 h post-transduction in lysis buffer. Use a minimum of 10,000 cells.
4. Incubate 4h - overnight at 55°C. After lysis, incubate the samples for 45 min at 85°C to inactivate the proteinase K.
5. Perform a standard (50 µL) PCR reaction using 2 µL of lysed cells (separate reactions of eVLP-targeted and control AECs) according to the manufacturer’s protocol.
6. Load 5 µL of PCR product onto a 1.5% agarose gel and perform agarose gel electrophoresis to verify that the PCR produced a single amplicon of the correct size (700 bp).
7. Perform a PCR clean-up and sequence the PCR fragment by Sanger sequencing with the forward PCR primer; see note 9.
8. Quantify editing efficiency using the ICE webtool by uploading the Sanger sequences [18]: [EditCo - CRISPR Performance Analysis](https://ice.editco.bio/#/).

**Figure 3:**
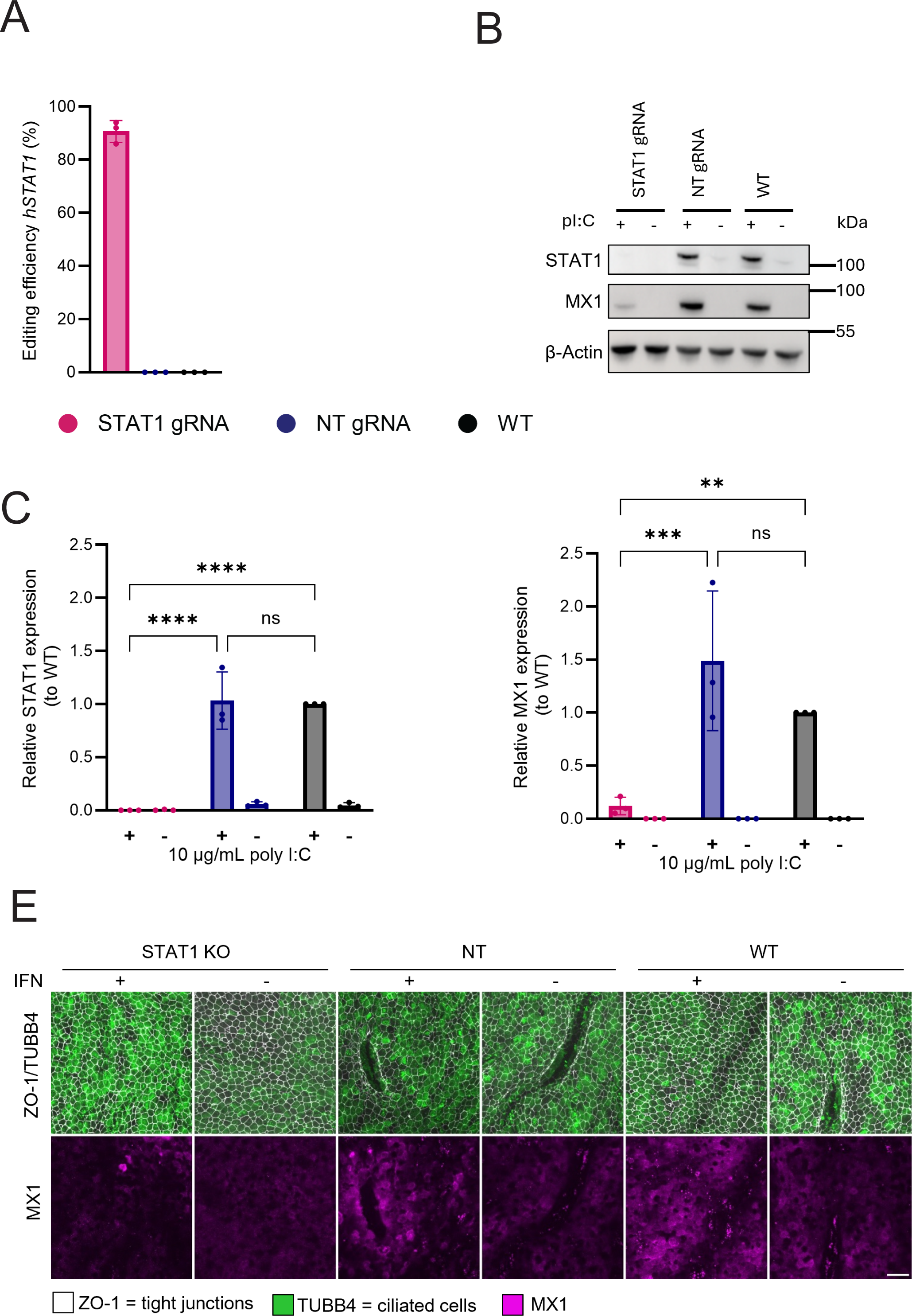
Efficient knockout of STAT1 in AECs using eVLPs. **(A)** Evaluation of gene editing efficiency by ICE analysis shows >90% editing of the STAT1 gene in AECs transduced with 0.2 μg/mL of eVLPs targeting *STAT1,* compared to AECs transduced with non-targeting (NT) eVLPs or untransduced, wild-type (WT) controls. N=3 independent biological replicates, bars represent the mean with standard deviation. gRNA sequence used for targeting STAT1 is 5’-GTCCACCAATGGCAGTCTGG-3’. **(B)** Western blot analysis of β-Actin, STAT1, and downstream MX1 expression shows efficient STAT1 depletion and reduced MX1 protein expression in AECs transduced with STAT1-targeting eVLPs, compared with NT or WT controls. Cells were treated with 10 μg/mL poly I:C (pI:C) (+) or left untreated (-) to induce ISG expression. Quantification of western blot protein expression of STAT1 **(C)** and MX1 **(D).** N=3 independent biological replicates, bars represent the mean with standard deviation. Ns p>0.05, * P ≤ 0.05, **P ≤ 0.01, *** P ≤ 0.001, **** ≤ 0.0001 by two-way Anova. **(E)** Maximum intensity Z-projections of well-differentiated STAT1 KO, NT, or WT ALI cultures show equivalent amounts of ciliated cells in STAT1 KO and control conditions, as well as reduced downstream MX1 expression in differentiated STAT1 KO cultures compared to controls. Inserts were differentiated for 6 weeks and stimulated with 50 ng/mL interferon lambda 3 (+) or left untreated (-) to induce ISG expression. They were subsequently fixed and stained with primary antibodies against ZO-1 (white), TUBB4 (green), and MX1 (magenta). The ZO-1 antibody was conjugated to Alexa fluor 594. For TUBB4 and MX1, the secondary antibodies donkey anti-rabbit Alexa fluor 488 and donkey-anti-mouse Alexa fluor 647 were used, respectively. bar = 20 μm.

### 3.3 KO AEC expansion and differentiation at ALI

Once KO AECs have been confirmed by ICE analysis or western blotting, they can be expanded for cryopreservation and subsequent subculture on porous supports.

#### 3.3.1 Organoid expansion of KO AEC

To expand KO AECs, cells are harvested after transduction and expanded as 3D airway organoids.

1. Prepare reagents by prewarming the cell culture plate at 37°C for a minimum of 1h and thawing 1 aliquot of BME on ice for 1-2 hours, or on ice in the fridge for 4 hours to overnight.
2. Visually inspect KO AECs 72 h post-transduction with a light microscope. Cells should be 70-80% confluent.
3. Collect medium from each well and wash the cell monolayer with HBSS.
4. Dissociate the cells with 250 µL/24-well of TrypLE™ express enzyme reagent by incubating the cell culture plate for 7-10 min in a humidified incubator at 37°C with 5% CO₂. Periodically check dissociation under an inverted light microscope.
5. Add the collected medium to the dissociated cells, pipet gently up and down to detach remaining cells from the wells and dissociate cells from clumps. Transfer to a 1.5 mL or 15 mL tube. Centrifuge at 500 x g for 5 min at RT.
6. Remove supernatant by pipetting without disturbing the cell pellet, resuspend in 1 mL HBSS to wash. Centrifuge at 500 x g for 5 min at RT.
7. Remove supernatant by pipetting and resuspend pellet in prewarmed pre-AOM. See note 10.
8. (Optional) Count cells with Trypan blue, see note 11.
9. Remove supernatant by pipetting and resuspend pellet very slowly in BME; avoid bubbles. Keep cells in BME on ice. See note 12.
10. Take your prewarmed plate out of the incubator. Add 30 µL of the BME/cell mix as a single dome to each well of a 48-well plate. Keep plates flat, and hold pipettes at a 90° angle to the plates. Start pipetting at the bottom of the plate and slowly move up while ejecting. Avoid bubbles.
11. Incubate the plate at room temperature for 1 min. Slowly turn your plate upside down and incubate in a humidified incubator at 37°C with 5% CO₂ for 10-15 min to polymerize the BME.
12. After incubation, turn the plate right-side up and add 300 µL of AOM to each well of a 48-well plate.
13. Return plates containing organoid cultures to the incubator. Incubate in a humidified incubator at 37°C with 5% CO₂ for 1-2 weeks until confluent (see Figure 4). Refresh AOM every 3-4 days.

**Figure 4:**
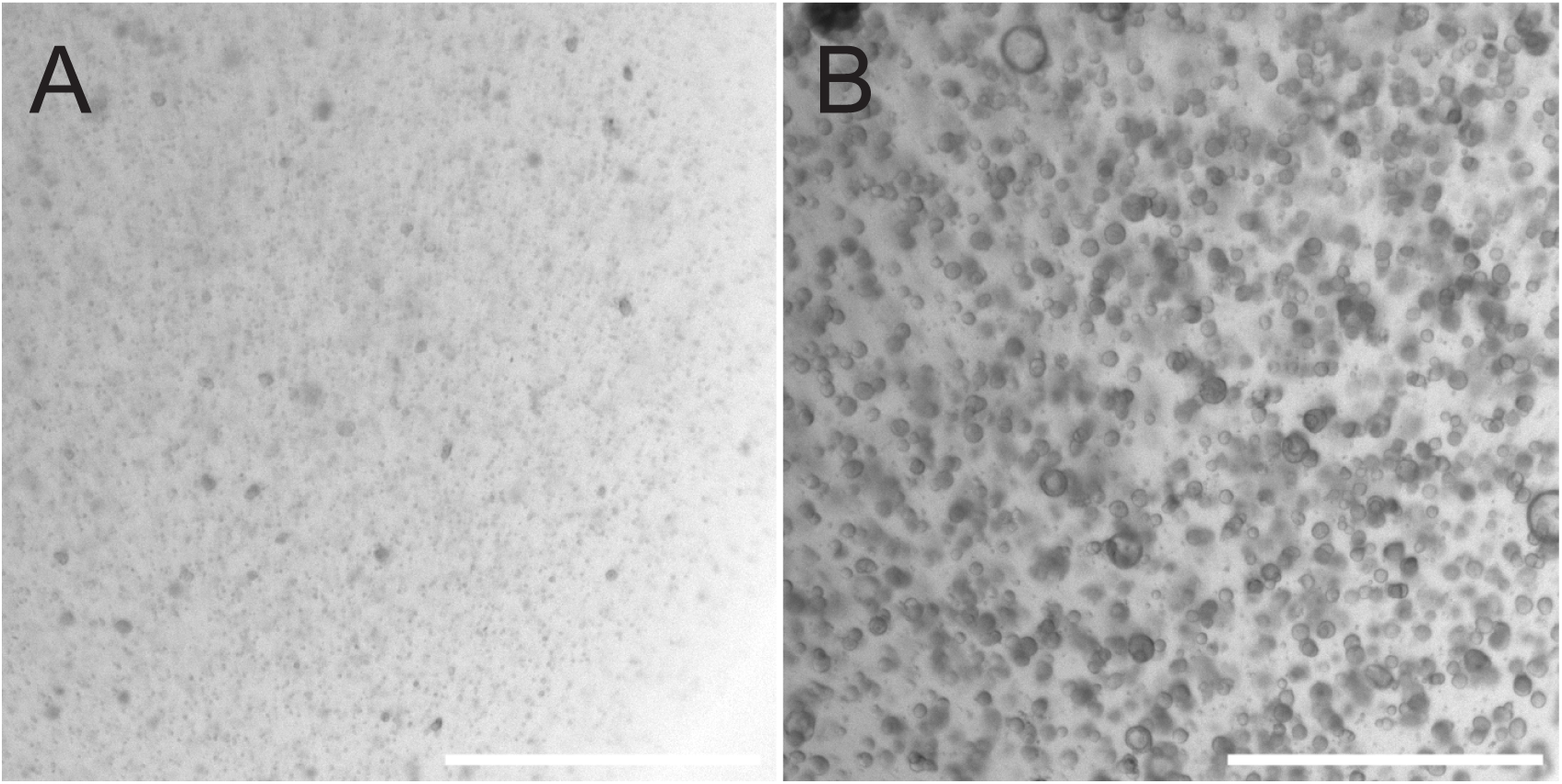
Expansion of AECs as organoids. Representative brightfield images of AECs cultured as organoids on day 0 **(A)** and day 12 **(B)** post-seeding. Cells were seeded with a seeding density of 1000 cells/µL. Scale bar = 1000 µm.

#### 3.3.2 Collagen Type IV Coating of Inserts

To ensure that AECs adhere properly to the insert membrane, the porous supports are first coated with collagen type IV.

1. Mix 9 volumes H_2_O with 1 volume Collagen Type IV. One 6.5 mm insert needs 50 µL of coating solution.
2. Pipet 50 µL into the apical compartment of one 24-well porous support chamber.
3. Tap the plate to ensure even distribution of the collagen solution.
4. Incubate the plates for 2-4 hours in a humidified incubator at 37°C with 5% CO₂.
5. Aspirate the coating solution, airdry for 5 min, and wash the apical compartment twice with PBS to remove traces of acetic acid.
6. Porous support chambers can be used directly or stored at 4°C for up to 6 weeks.

#### 3.3.3 Seeding, differentiation and maintenance of KO AECs

Following organoid expansion, KO AECs are seeded into collagen-coated porous support chambers and differentiated at the air-liquid interface for at least 3 weeks. The cells are seeded in EpiX medium to promote expansion in an initial liquid-liquid culture, until a confluent layer is established. Cells are then airlifted and cultured for 4-6 weeks in an air-liquid interface medium, which promotes mucociliary differentiation. As this process is lengthy, the cultures need to be maintained by replenishing medium and periodically removing mucus produced by airway secretory cells.

An example of differentiated knockout cultures can be found in Figure 3E.

1. Prewarm complete EpiX medium and add 500 µL basolaterally, 50 µL apically to each porous support chamber. Incubate in a humidified incubator at 37°C with 5% CO₂ during all subsequent incubation steps to prewarm and pre-humidify the porous support chambers.
2. Visually inspect your organoid domes. Domes should be confluent (see Figure 4) to isolate sufficient AECs to seed. See note 13.
3. Collect the old medium from the organoid domes.
4. Wash domes once with 300 µL HBSS per dome. Add 300 µL 1 mg/mL dispase solution supplemented with 5 µM Y-27632 to each well of a 48-well plate. Incubate in a humidified 37°C incubator for 2h.
5. Collect organoid domes with dispase in a 1.5-mL or 15-mL tube. Centrifuge for 5 min at 500 x g at RT. BME should be fully dissolved at this point.
6. Aspirate supernatant. Resuspend the pellet in TrypLE™ express reagent supplemented with 5 µM Y-27632 (approximately 0.1-0.2 mL per 30 µL dome). Incubate the tubes for 10-15 min at 37°C, pipet up and down every 5 min. Check periodically with a microscope whether organoids are fully dissociated.
7. When organoids are fully dissociated, add old medium collected in step 3 and gently pipet up and down to get rid of cell clumps and to inactivate the TrypLE™ express enzyme. Centrifuge for 5 min at 500 x g at RT.
8. Aspirate supernatant and resuspend pellet in 0.5-1 mL of pre-AOM supplemented with 5 µM Y-27632.
9. Count cells using trypan blue, then transfer to a fresh 1.5 mL or 15 mL tube. Centrifuge at 500 x g for 5 min at RT.
10. Aspirate supernatant and resuspend pellet in Epix medium with a density of 1x10^6 cells/mL. See note 14.
11. Add 100 µL of cells to the apical compartment of each porous support chamber, resulting in seeding density of 100,000 cells per insert. Incubate cultures in a humidified incubator at 37°C with 5% CO₂.
12. After 36-48h, change medium by gently aspirating the apical side, followed by the basolateral side. Add fresh EpiX medium to the basolateral side first, then to the apical side. Return cultures to a humidified incubator at 37°C with 5% CO₂.
13. Repeat medium change every 2-3 days with EpiX until confluence (see note 15).
14. Once cells are confluent, establish ALI by aspirating medium from the apical side and washing with 200 µL of TEER solution, added dropwise to the cells. Aspirate TEER solution, aspirate medium at the basolateral side and add 500 µL prewarmed ALI medium to the basolateral side only. Return cultures to a humidified incubator at 37°C with 5% CO₂. See also note 16.
15. After establishing ALI, maintain cultures by replenishing medium every 2-3 days to prevent acidification. Mucus (see note 17) on the apical side produced by the differentiating cultures can be removed by dropwise addition of 200-300 µL TEER solution, incubation of 10 min at 37°C, followed by gently pipetting up and down for at least 5x. Aspirate the remaining buffer.
16. Check the ALI cultures periodically to observe the stability of the cell layer and cilia formation. After 4 weeks of differentiation at ALI, depending on the donor, the cultures are ready for use. See also note 18.

#### 3.3.4 Cryopreserving KO AECs

Instead of direct differentiation at ALI, KO AECs can also be cryopreserved following steps 2-9 from section 3.3.3, frozen at -150°C and thawed when necessary.

1. Resuspend cell pellet in CryoStor CS10 with a density of 1,000,000 cells/mL. Aliquot cells in 250,000 cells/cryotube and keep on ice for 10 min.
2. Transfer the vials into a Corning® CoolCell™ and put the container at -80°C overnight, to allow for a controlled temperature decline.
3. The next day, move the cells to the -150° C freezer for long-term storage.
4. To thaw cryopreserved cells, dip a cryovial in a water bath (20-37°C) and transfer the contents to a 15 mL conical tube. Add 1 mL of prewarmed (37°C) pre-AOM, supplemented with CEPT, dropwise at a rate of one drop every 2 sec. Then slowly and dropwise add an additional 3 mL of prewarmed pre-AOM supplemented with CEPT at a rate of two drops every second. Mix slowly by pipetting up and down.
5. Centrifuge at 200 x g for 3 min at RT, aspirate supernatant, and resuspend pellet in 0.5 mL of prewarmed pre-AOM supplemented with CEPT.
6. Centrifuge at 200 x g for 3 min at RT. Aspirate supernatant and resuspend pellet in 180 µL ice-cold BME. Seed and culture organoids as described in steps 10-13 from section 3.3.1. Replace Y-27632 with 1x CEPT in AOM for seeding, continue culturing with regular AOM medium from the first medium change.

### 3.4 KO AEC infection

#### 3.4.1 Virus infection of KO AECs

1. Wash the apical side of the ALI cultures twice to remove mucus by adding 200-300 µL of TEER solution, incubating for 10 min at 37°C, then gently pipetting up and down at least 5 times. Aspirate the remaining buffer.
2. Refresh ALI medium on the basolateral side (see note 19). Return cultures to the incubator until use.
3. Prepare the inoculum by diluting virus to the desired titer in HBSS. Collect the inoculum and dilute 1:1 with 100 µL VTM and store at -80°C for later analysis.
4. Apply 100-200 µL inoculum at the apical side of the porous support chambers and incubate for 2 h in a humidified incubator at 33°C or 37°C with 5% CO₂ (see note 20).
5. Aspirate the inoculum, then wash the apical surface of the cultures three times with TEER solution by adding 200 µL of TEER solution dropwise and pipetting up and down 5x. Collect the third wash in an equal volume of VTM and store at -80°C for later analysis.
6. Return cultures to a humidified incubator at 33°C or 37°C with 5% CO₂ until the predetermined time points.
7. At each timepoint, collect viral progeny by adding 100 µL TEER solution dropwise to the apical side, incubating for 10 min at 33°C or 37°C, followed by gently pipetting up and down 5 times. Dilute apical wash 1:1 in VTM and store at -80°C for later analysis.
8. Cultures can be harvested for DNA, RNA, or protein analyses at any time point. Alternatively, cultures can be fixed and immunostained, as described previously [14].
9. Determine viral titer by tissue culture infectious dose (TCID_50_) or plaque assays, following established protocols (e.g. [13]).

## 4. Notes

1. Primary airway epithelial cells can be isolated directly from respiratory tract tissue, as previously described [14]. Alternatively, cells can be purchased commercially.
2. CEPT cocktail kit (Tocris) should be reconstituted according to the manufacturer’s protocol to the following stocks: Chroman 1 (0.5 mM, 10’000X), Emricasan (50 mM, 10’000X), trans-ISRIB (7 mM, 10’000X) and Polyamine (1’000X)
3. AOM can be prepared and stored at 4°C for 2 weeks, protected from light. Avoid freeze-thaw cycles of growth factors. Before use, bring AOM to room temperature.
4. A single gRNA is usually sufficient to create a >90% reduction of protein expression. To further increase editing efficiency, a combination of two (or more) gRNAs can also be used. Design a pair of gRNAs where the binding sites span an exon-intron junction of the target gene. The gRNAs should bind within a distance of 30-300 bp of each other.
5. As a control, select a gRNA that does not target the human genome (non-targeting, or NT), such as 5’-AAATGTGAGATCAGAGTAAT-3’. We recommend using NT eVLPs as controls throughout the protocol.
6. There is a large variety in the effectiveness of individual gRNAs. We therefore recommend testing 3-6 gRNAs per target gene before continuing with the protocol. For initial experiments, we recommend preparing 2 mL eVLP stocks of each gRNA, skipping the concentration (3.1.3) and quantification (3.1.4) step and directly transducing either AECs or cell lines such as A549 or BCI-NS1.1 with undiluted eVLPs by following the steps in 3.2.1-3.2.3. Efficacy of the gRNAs can be evaluated as described in part 3.2.4. Efficient gRNAs should reach 80-90% editing efficiency in AECs using unconcentrated eVLPs.
7. Repeated concentrations of bigger volumes take longer than initial concentrations.
8. For efficient gRNA delivery, a concentration of 0.2-0.4 µg/mL Cas9 is sufficient and well tolerated by AECs. For less efficient gRNAs, a higher dose could be tried, but cell health should be carefully monitored.
9. For ICE analysis, make sure Sanger sequence traces are untrimmed after sequencing.
10. Optionally, add 5 µM Y-27632 to pre-AOM to prevent cell death. This is recommended when passaging multiple KO lines simultaneously.
11. Small numbers of cells are difficult to count. For transduction in a 24-well plate, assume a cell density of around 250,000 cells 72 h post-transduction.
12. Aim for 1000-1500 cells/ µL BME. For cells transduced in a 24-well, we recommend resuspending the cell pellet in 180 µL BME, creating 6 domes. If it is difficult to pipet off all the remaining supernatant, the BME can be diluted to 70% in the cell mixture without significantly affecting organoid growth.
13. One full dome of organoids yields approximately 100,000-250,000 cells after 1-1.5 weeks of culture, depending on the donor and the cells’ viability before seeding.
14. Alternatively, cells can be further expanded into organoids by resuspending the cell pellet in BME (1000-1500 cells/µL) and following steps 9-13 of section 3.3.1.
15. Cells can generally be airlifted after 4-7 days of expansion. Airlifting too early, when the epithelium is not yet confluent enough, will lead to leakage. Airlifting too late can also have detrimental effects on epithelial health, and cells expanded for longer than 7 days on inserts usually fail to form well-differentiated cultures. Make sure the epithelium has a consistent cobblestone architecture throughout the insert (no stretched cells remaining) for at least 1 day before airlifting.
16. Check the cells the day after establishing ALI. If medium leaks from the basolateral side to the apical surface, add 200 µL of ALI medium to the apical side and repeat ALI establishment within 1 or 2 days.
17. Mucus protects the cells but can accumulate over time and negatively affect them. Therefore, wash the apical surface with TEER solution only when mucus clearly accumulates.
18. If cells deteriorate at ALI or if the epithelium is scratched during handling, boost cell proliferation and epithelial recovery by adding a gradient of epidermal growth factor (EGF) to the ALI medium. Start with a high dose of EGF (25 ng/mL, 1:1000) for the first medium change, followed by 10 ng/mL, 5 ng/mL, and 2.5 ng/mL in the following medium changes. EGF stocks can be prepared by dissolving 1 mg of long EGF (Repligen) in 1 mL of 10 mM HCl solution. Dilute in 39 mL PBS to 25 µg/mL and filter-sterilize through a 0.22 µm filter. Aliquot and store at -20°C for up to 1 year. Alternatively, use a 2:3, 1:4 and 1:9 ratio of EpiX medium with ALI medium for the subsequent 3 medium changes.
19. For some viruses, infection can be boosted by removing heparin and hydrocortisone 2-3 days before and during infection. This should be experimentally determined for each virus.
20. Optimal culture temperature should be experimentally determined for each virus.

## 5. Expected results

This workflow produces genetically modified primary airway epithelial cells that can be expanded as organoids and then differentiated at the air–liquid interface. Figures 2 and 4 show what these cells look like at key stages: before and after eVLP delivery (Figure 2) and during organoid expansion (Figure 4). Figure 3 demonstrates the full pipeline using STAT1 knockout as a test case.

In this example, genomic analysis showed >90% editing at the STAT1 locus in primary AECs treated with STAT1-targeting eVLPs (Figure 3A). Western blot confirmed that STAT1 protein was depleted and that downstream MX1 expression was reduced after stimulation, relative to non-targeting and untransduced controls (Figure 3B–D).

After expansion and differentiation at the air–liquid interface, immunofluorescence images showed ciliated cells in both STAT1-targeted and control cultures, with reduced MX1 staining in stimulated STAT1-targeted cultures (Figure 3E). Importantly, the edited cells retained the ability to form differentiated airway epithelium.

Taken together, these examples demonstrate what you can expect from the workflow: efficient editing at the chosen locus, corresponding protein depletion, and differentiation into functional airway cultures with the expected phenotype. The STAT1 case illustrates the approach; actual editing efficiency and effects on expansion or differentiation will vary depending on the target gene and the donor cells used.

## Acknowledgments

This research was supported by the Swiss National Science Foundation (Grant No. 310030_207675). We would like to express our thanks to Dan Wettstein for his support in, image acquisition, and processing. Furthermore, we are grateful to Sabina Berezowska and Irene Ramos-Centeno (Institute of Pathology, University of Bern) for providing the respiratory tract tissues via the Tissue Bank Bern. Microscopic imaging was performed on equipment supported by the Microscopy Imaging Center (MIC), University of Bern, Switzerland

